# Unintentional finger force drifts are minimally influenced by temporal evolution of surface friction

**DOI:** 10.64898/2026.06.25.734530

**Authors:** Mia Naranjo, Sophia Rockland, Sasha Reschechtko

## Abstract

Humans consistently decrease the amount of force they produce during isometric finger pressing in the absence of visual feedback, a phenomenon often called “force drift.” This decrease in force production has been attributed to limitations in working memory and/or adaptive neural control processes that minimize energy consumption. In this study, we investigated a potential peripheral reason for such force drifts: increases in the coefficient of friction between the fingertip and the surface it contacts due to changes in fingertip contact area as the fingertip hydrates under prolonged pressure. We investigated this possibility by eliciting force drifts from participants performing isometric pressing tasks against smooth glass, which shows the phenomenon of increasing contact area during prolonged contact, and a polymer which does not exhibit this phenomenon. We confirmed that the coefficient of friction only increased on the glass plate, however we did not observe a difference in force drifts between these two surfaces, although we found some evidence that force drift could be associated with coefficient of friction. Our findings suggest that factors other than peripheral changes in coefficient of friction are the primary drivers of force drifts.

## Introduction

The ability to pick up and hold objects is an integral part of many activities of daily living. While maintaining sufficient grip force is necessary to keep an object stable in hand, unintentional decreases in finger force production have been well-documented. Humans typically have difficulty maintaining constant force production in isometric conditions where the fingers press against an immovable surface. Vaillancourt and Russel (2002) showed that finger force production consistently in isometric conditions consistently decreases once it exceeds a relatively low force level. This decrease, or drift, in force production is apparently involuntary: it occurs when participants are asked to continue to press with the same amount of force, and participants appear unaware of them (Cuadra and Latash, 2019; Jo et al., 2016; Reschechtko et al., 2017a). Unintentional decreases in finger force production have been explained by some groups as a limitation in short-term visuomotor memory (Coombes et al., 2011; Poon et al., 2012; Vaillancourt et al., 2003; Vaillancourt and Russell, 2002). Force drifts have also been taken as an example of “motor slacking” (Reinkensmeyer et al., 2009; Smith et al., 2018), or the tendency to reduce muscular effort and explained as a tendency of natural systems to achieve states of lower potential energy, brought about by changes in control variables associated with force production (Abolins et al., 2023; Abolins and Latash, 2022; Ambike et al., 2016).

While various factors have been proposed to account for decreases in finger force production, such decrease seems potentially detrimental to everyday tasks involving object manipulation. To hold an object in hand, the grip force applied to the object must be sufficient to maintain tangential forces – often called “load forces” in the context of object manipulation – greater than weight of the object being held. The relationship between grip and load force depends on the coefficient of friction between the fingers and surface, which in turn depends on the surface conditions. Object manipulation forces are very sensitive to changes in surface texture (Johansson and Westling, 1984) and can rapidly change as small slips between the fingertip and object are detected (Barrea et al., 2018; Delhaye et al., 2021a; Johansson and Westling, 1988, 1987). Despite these mechanisms for addressing object slips, there is evidence that force drifts occur in situations where objects are held in hand (Naik and Ambike, 2022; Smith et al., 2018) and not just during isometric conditions where force production is less related to manipulation forces. However, given the close relationship between finger force production and object manipulation forces, interactions between force drifts and surface conditions – particularly those affecting the coefficient of friction – seems plausible.

A major factor affecting coefficient of friction is the contact area between the fingertip and the surface it contacts. Although many studies have implicitly assumed this contact area to be constant, in many cases it actually evolves over time (Dzidek et al., 2017, 2016; Pasumarty et al., 2011). This evolution of contact area occurs when the fingertip is in prolonged contact with a nonporous surface, causing it to become “occluded” or hydrated by sweat. This process softens the fingerprint ridges and allows them to conform more completely with the surface. Fingerprint ridges are part of the most superficial layer of skin, stratum corneum, which contains a large proportion of keratin. This keratin contains both crystalline and amorphous molecules; when they become wet (by hydration through occlusion), the amorphous molecules become much more elastic. Therefore, when fingertips are in contact with a surface harder than keratin, hydration causes the contact area between the fingertip and underlying surface to increase, which subsequently increases the coefficient of friction. In contrast, when the fingers are in contact with a surface softer than keratin, the surface conforms to fingerprint ridges almost immediately and contact area changes minimally over time.

This hydration process (evolution of friction) and force drifts both occur over tens of seconds; this similarity suggests the possibility that these processes are related. In particular, it raises the possibility that force drifts are an adaptive process used by the nervous system to minimize energy expenditure as fingertip friction increases. We carried out this study to investigate this possibility by measuring fingertip coefficient of friction and unintentional force drifts. Our experiment involved a standard isometric force production task that led to force drifts (Ambike et al., 2016; Reschechtko et al., 2017b; Vaillancourt and Russell, 2002) and varied the surfaces that participants pressed against. In some trials, participants pressed against smooth glass, which elicited changes in coefficient of friction over time. In other trials, participants pressed against glass which had been coated in polydimethylsiloxane (PDMS), a silicone polymer softer than keratin which therefore does not elicit changes in temporal evolution of frictional force. If force drifts were indeed related to changes in the coefficient of friction, we expected to see (1) that unintentional force drifts will occur at different rates on these two surfaces and (2) that there was a relatively consistent relationship between the magnitude of force drifts and level of fingertip friction. While we found evidence of the second hypothesis, we did not find evidence for the first.

## Methods

### Participants

20 healthy young adults (ages 18-35; 11 women) participated in the study. Participants inclusion criteria were: age between 18-45 years, had the ability to sit and complete data collection for a 90-minute period, and no history of injuries to their dominant hand that would inhibit use of the fingers. All participants were right-handed. All participants performed experimental procedures approved by the San Diego State University Institutional Review Board’s Human Research Protection Program. Participants received $25 USD compensation for their participation.

### Materials

Finger force production was recorded with a multi-axis force and torque sensor (Axia-80, ATI Industrial Automation, Apex NC). A 3D-printed platform supporting two glass plates (dimensions) was fixed to the sensor. One of the glass plates was covered in a layer of a silicone elastomer, polydimethylsiloxane (PDMS; Sylgard-184, Dow Inc., Midland, MI), while the second glass plate was left untreated and smooth. Participants received visual feedback about the amount of downward (normal) force they applied to the glass plate on a 27” computer monitor. The feedback consisted of a vertical “tank” that filled in proportion to force applied; the target force people were asked to maintain was set in the center of the bar. A program written in the LabVIEW programming environment (Version 2022; NI, Austin, TX) was used to provide participants with visual feedback on the forces they applied and log force data for offline analysis. Schematics of the experimental setup and participant feedback are presented in Figure 1 panelsA-C.

**Figure 1:**
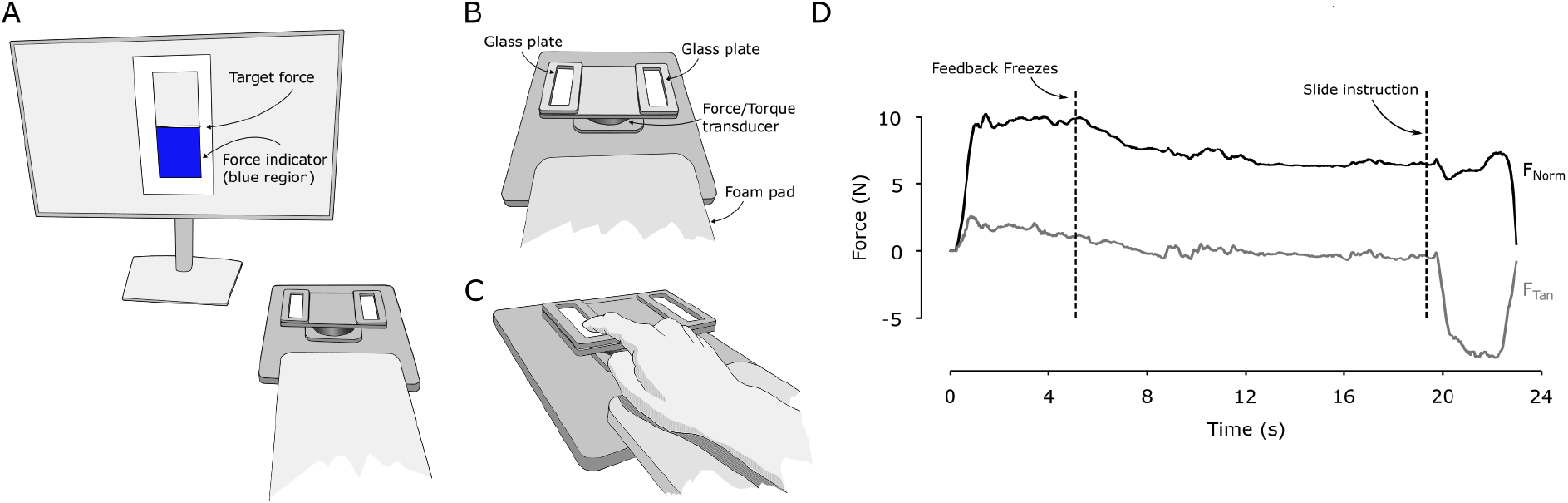
Experimental setup and example trial. Panel A: Experimental setup with monitor and force transducer. Participants pressed on the force transducer (refer to Panels B and C) to fill the target indicator to the halfway point. Panel B: Schematic of the force transducer with attachment to hold two glass plates. Either the left or right plate had a PDMS coated glass slide, while the other one had a smooth glass plate. Panel C: Schematic showing how participants interacted with the glass slides, resting their forearms on a foam pad and pressing on the indicated plate with their index fingers only. Panel D: Example force profiles for normal force and tangential force during an experimental trial. Participants received feedback (see Panel A) on the normal forces they produced.

### Procedures

A member of the study team oriented participants to the set up after they reviewed and signed a consent form. Before beginning the experimental procedure, the participants washed and dried their hands to normalize skin conditions. During the orientation, participants pressed on a glass plate to complete practice trials of the procedure with the target force set at either 5 or 10 Newtons (N) for 20 seconds. Participants completed practice trials until they were able to quickly meet the goal force level and were confident in their ability to maintain that instructed force level.

During experimental trials, participants were instructed to press on the plate to reach the goal force level, which was always 10 N. Veridical visual feedback was provided during this time as a “tank” that filled according to the level of force produced (Figure 1A). After a random interval between 4 and 7 seconds, the visual feedback “froze” at the target level such that, even if participants changed the amount of force they were producing, the visual feedback they received did not change. Participants were not informed when or that this change occurred. Finally, 2-20 seconds after the feedback froze, the force feedback changed color to indicate to the participant that they should slide their finger across the plate (toward their body) and then release; each trial ended when the participant lifted their finger from the glass plate. Force traces and instructions for an example trial are provided in Figure 1D. Each participant completed 60 trials and switched between the plates in a quasi-random order such that they performed 30 trials of varying durations on each plate. Both the amount of time before feedback froze (4-7 seconds) and the duration of the trial after visual feedback froze (2-20 seconds) were selected quasi-randomly from a flat distribution.

### Kinetics

We recorded the forces and torques that participants applied using a 6-axis force/torque transducer (Axia-80, ATI Industrial Automation, Apex NC, USA) sampling at 500 Hz.The glass plates (one bare and one covered in PDMS) were rigidly mounted to the transducer with 3D printed fixtures.

### Analysis

We imposed some trial acceptance criteria to ensure that data analyzed were indicative of the instructed task. For each trial, we checked whether the participant’s normal force was within 1.5 N of the instructed force level in a time window from 25 ms before feedback froze to 25 ms after feedback froze. Trials in which normal force did not fall within this tolerance range were discarded. For trials with acceptable normal forces, we also double-checked in make sure that participants were pressing on the correct surface by computing the point of force application using the formula d_x_ = τ_x_/F_z_ where d_x_ is the x coordinate of force application, τ_x_ is the torque applied about the x axis, and F_z_ is the normal force. If a participant was found to have been pressing on the other surface for a given trial, that trial was reclassified as the other surface condition for analysis. The maximum number of trials excluded from a surface condition according to these criteria for an individual participant was 6 (of 30), and usually 3 or fewer trials were excluded from a condition.

Due to the experimental design, each participant performed the same number of trials per surface condition, but these trials were not the same duration as each other. This is because the duration of each trial varied randomly to allow us to investigate the evolution of frictional force over time. Therefore, we used two methods to compare the speed of force drifts between conditions and test the hypothesis that force drifts would vary depending on the surface participants pressed against. One method investigated short-term force drifts across trials by averaging each participant’s force production for the first two seconds after force feedback was frozen; this time period included data from all of the accepted trials because they all lasted longer than 2 seconds; we then used those data to fit an exponential function of the form F(t) = a ✕e^(-t/τ)^ + c where F(t) is the normal force at time t, a and c are coefficients representing the net change as t → ∞, and τ is a time constant describing the speed of change. We subsequently compared values of τ between each surface using a paired t-test (scipy’s *ttest_rel* function).Our other method binned trial duration into 2 second bins and averaged the final force level of each trial within that bin to compute a low-resolution estimate of force drift over time. We analyzed the level of force drift over time using a two-way repeated measures ANOVA with factors *time* (9 levels, one for each 2-s time bin) and *surface* (two levels, for PDMS-coated and uncoated glass).

We also approximated the coefficient of static friction (μ_s_) that resulted from the combination of force drift and fingerprint conformity in each trial. μ_s_ was computed as the ratio of the tangential force to the normal force, at a 20 ms time window surrounding the time when the fingertip began to slip. We identified the onset of slip by finding when the tangential force began to decrease following its initial increase after the participant was instructed to drag their finger across the surface. In order to verify that coefficients of friction were different between the two surfaces (to ensure that our experimental manipulation was working), we binned the μ_s_ according to the time when they occurred and used the two-way repeated-measures ANOVA time x surface to investigate force drift over time. To test our second hypothesis – that force drift is related to coefficient of friction – we investigated whether there was any systematic relationship between force drift and coefficient of friction using repeated measures correlation that took into account each accepted trial from each participant.

With the exception of repeated measures correlation and t-tests on short-term force drifts, we performed statistical analyses in JASP (JASP Team, 2024). We performed repeated measures correlation in Python as implemented in the Pingouin package (Vallat, 2018). Unless otherwise noted, descriptive statistics are presented as mean ± standard deviation.

## Results

Participants generally completed the testing in less than 1 hour. Of the 20 participants tested, all but one demonstrated robust force drifts when force feedback was frozen.

### Evolution of frictional forces differed depending on surface

We verified that our experimental manipulation had the intended effect on fingertip friction by investigating whether the coefficient of friction increased in a time dependent manner on either surface. Our expectation was that, on the smooth glass surface, coefficient of friction would increase as the duration of the trial increased, due to an increase in real contact area between the fingertip and surface. In contrast, we did not expect that coefficient of friction would be affected by trial duration on the PDMS-coated surface because the compliant PDMS conforms to fingerprint ridges. We tested whether the coefficient of friction consistently changed over time on each surface by running a repeated measures correlation on coefficient of friction and trial duration for all trials each participant performed on a given surface. For the smooth glass surface, there was a positive correlation between trial duration and the coefficient of friction (r = 0.213; P = 7.007e^-7^) indicating a consistent increase in coefficient of friction as the contact time increased (Figure 2A). In contrast, for the PDMS-coated surface, there was no correlation between time and coefficient of friction (r = 0.002; P = 0.952; Figure 2B).

**Figure 2:**
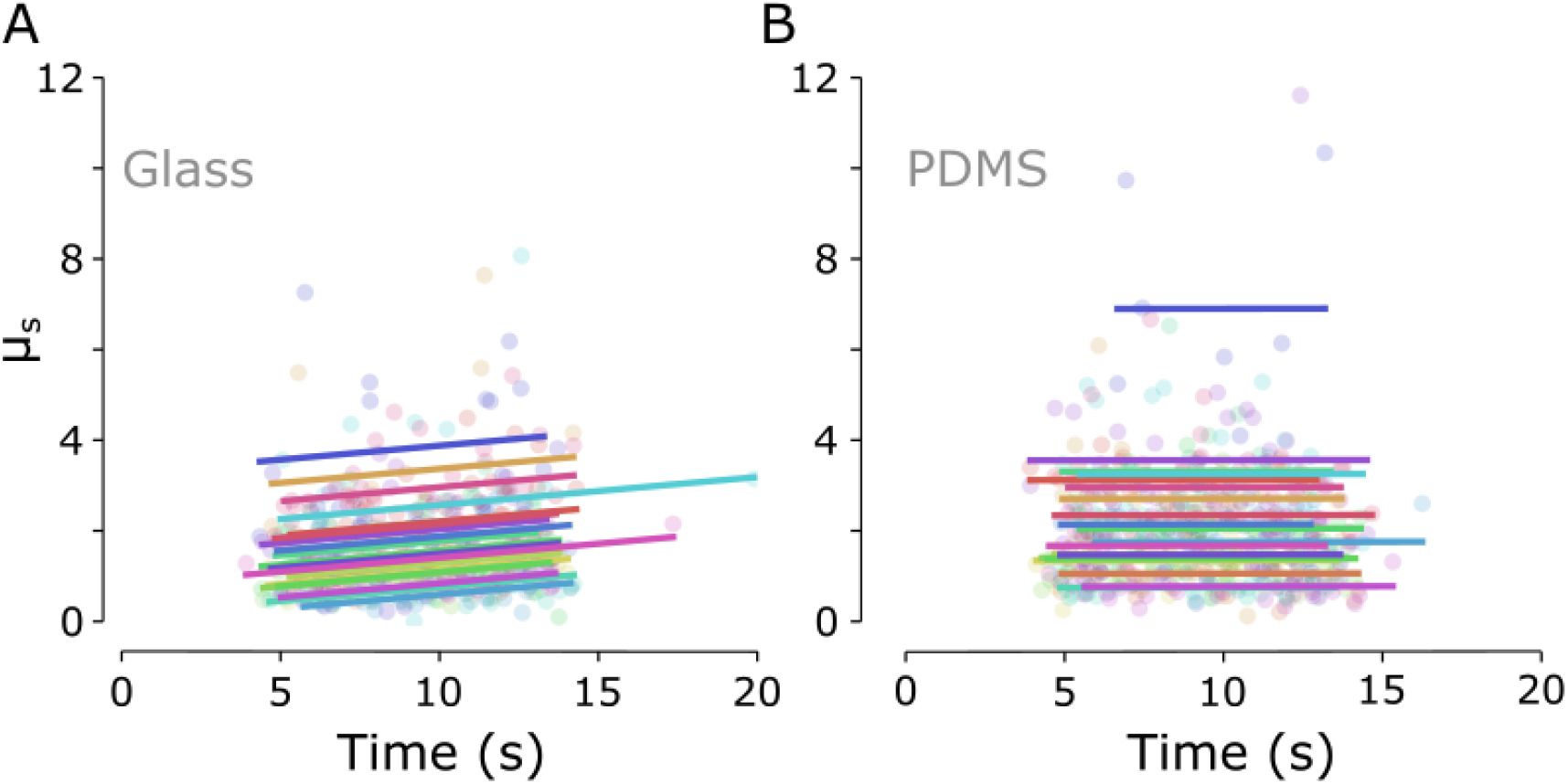
temporal evolution of static friction on different surfaces. Panels A and B plot each accepted trial for each participant showing the time elapsed since trial onset (x axis) and computed coefficient of friction (μ_s_) at the end of the trial (y axis). Panel A shows these data when participants pressed against a clean glass surface; Panel B shows these data when the same participants pressed against a glass surface coated in PDMS polymer. Each color represents a single participant and each dot represents a single accepted trial. Lines illustrate best fits for each individual participant; note the consistent positive slope in panel A compared to negligible slope in Panel B.

### Short-term decreases in force production were unaffected by surface

Force drifts typically begin very soon after the loss of visual feedback. All trials lasted at least 2 seconds after visual feedback was frozen, so we used these data from all trials to investigate the initial decrease in force production. For each participant, we compared the time constants for fitted decays between the two surfaces using a paired t-test. For the PDMS coated surface, the time constant was 2.47 ± 1.77 s, whereas it was 3.97 ± 5.46 s for the smooth glass, without evidence of a consistent difference between the fitted time constants (T_19_ = 1.298; P = 0.210).

### No consistent difference in force drift between surfaces

The duration of each individual trial was quasi-randomly assigned so that each participant performed 30 trials per surface with durations between 2 and 20 seconds. For each participant, we binned the trials into 2 second bins according to duration. For each bin, we then found the median change in force (final force level – initial force level) for all trials that fell into that bin for each surface condition. We used a linear mixed effects model with fixed factors *bin* (1-9 representing 2 second bins of trial duration following feedback freezing) and *surface* (PDMS and glass). Consistent with the visual impression from Figure 3, there was a significant effect of bin (F_8,32.32_ = 8.624; P = 3.273e^-6^) – reflecting the existence of force drifts – but neither *surface* nor the *bin*✕*surface* interaction were significant (F_1,25.55_ = 0.199; P = 0.659 and F_8,236.56_ = 1.644; P = 0.113, respectively).

**Figure 3:**
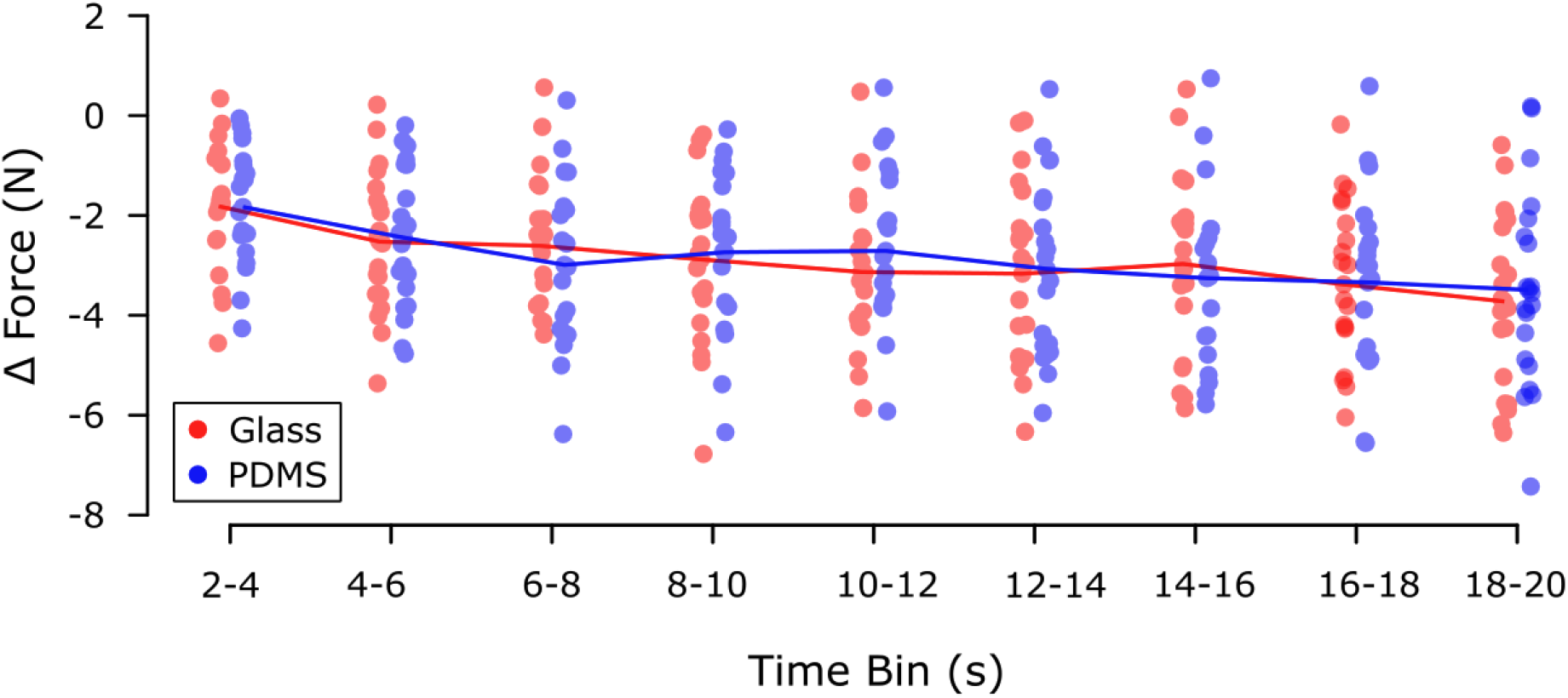
force drifts over time. Average change in normal force (y axis) plotted against the time bin for which the average was computed. Each point represents a single participant’s average force change in that time bin. Blue points represent averages when participants pressed against a smooth glass plate; red points represent averages when participants pressed against PDMS coated glass.

### Coefficient of friction was associated with the magnitude of force drift

The main hypothesis tested in this study was that force drifts are related to time-dependent changes in the coefficient of static friction. In particular, we wanted to investigate whether force drifts take coefficient of friction into account, such that larger coefficients of friction are associated with larger decreases in force. To test this, we conducted a repeated measures correlation analysis to investigate whether there was any association between force drift and coefficient of friction on a trial-by-trial basis. Repeated measures correlation showed strong positive associations between coefficient of friction and force drift for both smooth glass (r = 0.435; P = 4.450e^-26^) and PDMS (r = 0.477; P = 4.591e^-31^). The sign of the correlation coefficient indicates larger decreases in force were observed on trials where coefficient of friction was larger, regardless of whether the participants were pressing against smoothg glass (Figure 4A) or PDMS coated glass (Figure 4B).

**Figure 4:**
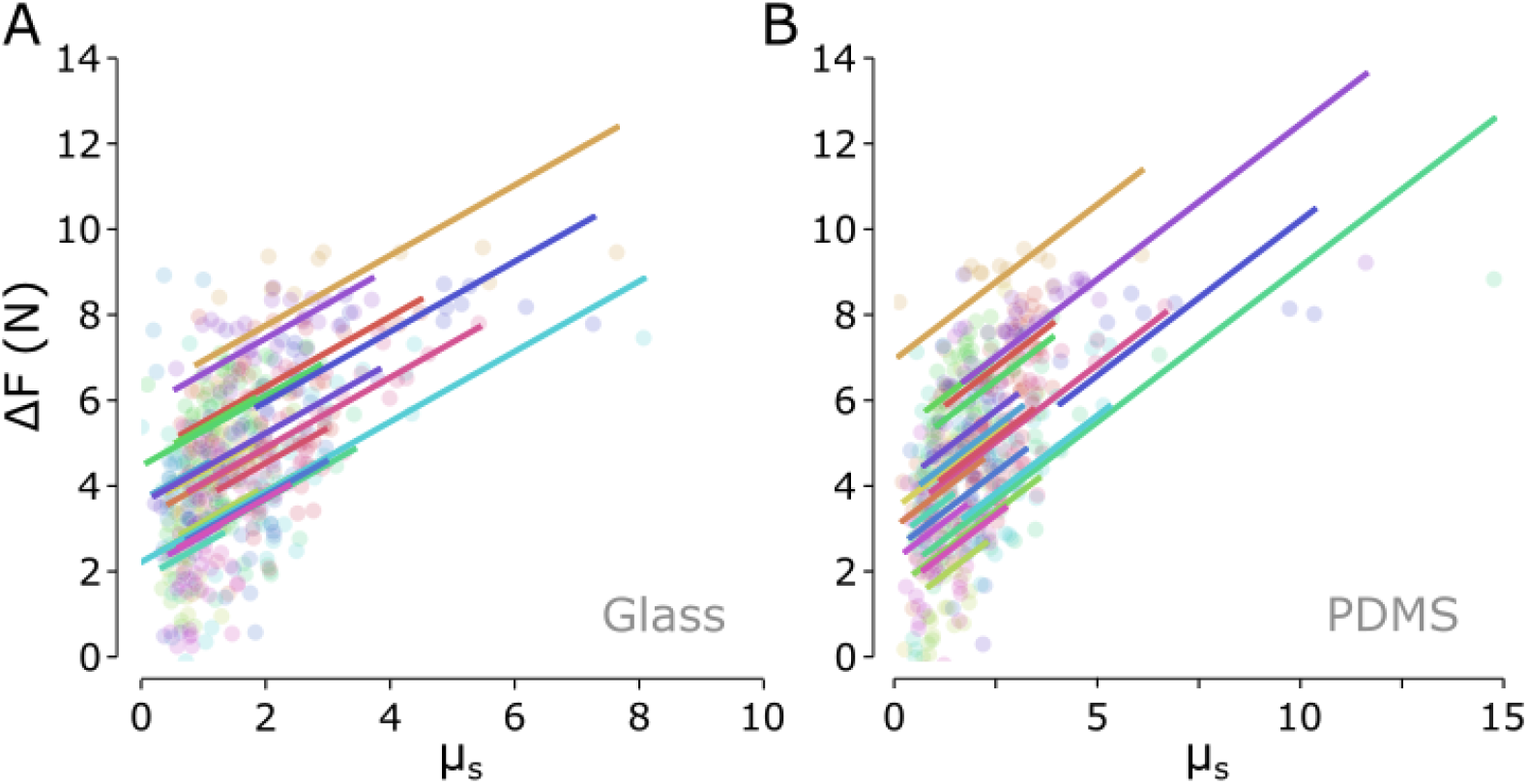
association between static coefficient of friction and force drift. Panels A and B plot each accepted trial for each participant showing the computed static coefficient of friction (μ_s_) at the end of the trial (x axis) and force drift since feedback freeze (y axis). Panel A shows these data when participants pressed against a clean glass surface; Panel B shows these data when the same participants pressed against a glass surface coated in PDMS polymer. Each color represents a single participant and each dot represents a single accepted trial. Lines illustrate best fits for each individual participant.

## Discussion

We performed this study to investigate the possibility that the well-documented phenomenon of isometric force drifts (Ambike et al., 2016; Reschechtko et al., 2017a; Vaillancourt and Russell, 2002) could be explained by the evolution of fingertip contact area (Dzidek et al., 2017, 2016, 2014), and therefore coefficient of friction, during prolonged object contact. We tested two primary hypotheses: first, that unintentional force drifts occur at different rates on different surfaces (depending on whether they allow for temporal evolution of friction), and second that magnitudes of force drift are associated with fingertip friction. Our findings do not support the first hypothesis but are consistent with the second one. Overall, our observations provide equivocal support for fingertip friction as a contributing factor, but are not consistent with within-trial evolution of fingertip friction playing a major role in fingertip force drifts. In addition to the hypothesis tested, our study provides additional insights into the tribology of the fingertip under more ecological conditions than have been previously studied.

### Minimal evidence for force drift sensitivity to frictional forces

Our experimental manipulation successfully elicited changes in evolution of fingertip friction between clean glass and PDMS surfaces. This change is shown by strong correlation between time and friction on clean glass contrasted by no correlation between time and friction on the PDMS coated glass. Nonetheless, force drifts were not appreciably different when participants pressed against the two surfaces investigated, which show qualitatively different characteristics of evolution of friction. This does not support the hypothesis that changes in fingertip friction is a primary driver for fingertip force drifts.

The strongest evidence for a relationship between force drift and coefficient of friction comes from the strong correlations observed between the two measures across both surfaces, which comes with some computational caveats. In particular, the coefficient of friction was computed from normal force, so they will necessarily co-vary. In an effort to account for this source of covariation, we did not directly correlate normal forces with the computed coefficient of friction but rather with change in force after feedback was frozen. Because participants do not have perfect performance, they start from different force levels initially and change in force will not have a 1-to-1 correspondence with the normal force levels used to compute coefficient of friction.

### Proposed mechanisms of force drift

If the evolution of fingertip friction is not a major player in force drifts, what mechanisms might be? Broadly, different researchers have proposed two categories of mechanisms to account for these force drifts. Some researchers have focused on force production without visual feedback as “memory guided” force production, and therefore interpret force drifts as the result of limitations in memory (Coombes et al., 2011; Vaillancourt et al., 2003; Vaillancourt and Russell, 2002). Support for such an interpretation comes from imaging studies (Poon et al., 2012; Vaillancourt et al., 2003) as well as the observation of changed force drift characteristics in populations with neurological disorders or under scenarios such as sleep deprivation (Brinkerhoff et al., 2022; Neely et al., 2019; Vaillancourt et al., 2001).

On the other hand, some experimental findings are not obviously compatible with force drifts as a result of memory limitations. One question is why people would always “forget” in the same direction – i.e. toward force decreases. Similarly, force drifts seem to show sensitivity to force production level, such that larger drifts are observed when starting from higher force production levels no force drifts are observed and at some very low levels of force (Ambike et al., 2016; Jo et al., 2016; Naik and Ambike, 2022; Vaillancourt and Russell, 2002); it is not clear why a higher level of force would be less “memorable.” Finally, these force drifts seem to occur during continuous force production, but are not present when participants perform pressing after a delay (Cuadra and Latash, 2019; Jo et al., 2016; Reschechtko et al., 2017a), again casting doubt on memory – which would degrade over time – as a primary explanation. An alternative explanation of involuntary force drifts has been proposed that they are adaptive strategies related to a tendency for the motor system to favor strategies that decrease exertion in the absence of an error signal. This idea is sometimes called “motor slacking” (Reinkensmeyer et al., 2009; Smith et al., 2018) and it is broadly compatible with observations of minimal intervention in motor control including optimal feedback control (Diedrichsen et al., 2010; Todorov and Jordan, 2002) and the uncontrolled manifold hypothesis (Scholz et al., 2000). Our study does not provide straightforward evidence for either mechanism of force drift, although changes in force production related to friction seem more compatible with the motor slacking hypothesis.

### Occlusion-driven friction changes occur at grasp-relevant force levels

Dzizek and colleagues (2017) previously showed the temporal evolution of fingertip friction during contact with glass surfaces. Their study showed the phenomenon in two human participants at servo-controlled force levels of 1.5 N. In contrast, the present study extends the observation of this phenomenon to more ecological settings in 20 human participants. Participants in our study used self-generated forces to press against a force sensor at levels about an order of magnitude higher (target level of 10 N).

While forces around or even under 1 N (or lower) are common in tasks that require tactile acuity such as search or identification (Olczak et al., 2018; Peters et al., 2009; Reschechtko et al., 2024), higher levels of force production are more common in object manipulation, which is a common scenario in which participants might be in contact with an object for an extended period of time. Holding an object for an extended period of time – such as holding a glass at a social function – is a more likely time when changes in surface friction could affect behavior. As such, one of this study’s most important contributions is the extension of Dzizek and colleagues’ findings to levels of force more associated with scenarios where the evolution of fingertip friction could be ecologically relevant.

### Limitations

We carried out this study to investigate changes in fingertip coefficient of friction as a peripheral mechanism affecting unintentional finger force drifts. One primary difficulty in assessing the association between drifts and coefficient of friction is the computational relationship between the measures since coefficient of friction is computed relative to the normal force. One way of disentangling these factors in the future could be investigating drifts in tangential finger forces. Tangential finger forces can be controlled relatively independently of normal forces (Reschechtko et al., 2017b) and coefficient of friction is not computed directly from it. Another possibility would be measuring the evolution of the true fingertip contact area (Bochereau et al., 2017; Delhaye et al., 2021b) rather than coefficient of friction. This would allow measurement of a factor analogous to coefficient of friction without requiring participants to make kinetic adjustments (e.g. sliding their fingers at the end of the trial) that might introduce other control factors into observations.

Related to this previous point, the computation of coefficient of friction in our study required that participants slide their fingers across a surface at the end of each trial. This may have introduced a secondary goal for participants (other than maintenance of fingertip force), which could have unintended consequences for force production. Participants were not aware of when each trial would conclude, so while they were not able to anticipate this occurrence, the knowledge that they would have to do something else later could have affected their performance (Naik and Ambike, 2022). Having trials of varying and randomized lengths also affected the precision of our force drift estimates because we needed to bin data. While we do not think that this seriously degraded the veracity of our force drift measures, simply measuring force drifts over tens of seconds on surfaces with different coefficients of friction could provide a simple analysis to verify the results we observed.

## References

Abolins, V., Latash, M.L., 2022. Unintentional force drifts across the human fingers: implications for the neural control of finger tasks. Exp. Brain Res. 240, 751–761. 10.1007/s00221-021-06287-2

Abolins, V., Ormanis, J., Latash, M.L., 2023. Unintentional drifts in performance during one-hand and two-hand finger force production. Exp. Brain Res. 10.1007/s00221-023-06559-z

Ambike, S., Mattos, D., Zatsiorsky, V.M., Latash, M.L., 2016. Unsteady steady-states: central causes of unintentional force drift. Exp. Brain Res. 234, 3597–3611. 10.1007/s00221-016-4757-7

Barrea, A., Delhaye, B.P., Lefèvre, P., Thonnard, J.-L., 2018. Perception of partial slips under tangential loading of the fingertip. Sci. Rep. 8, 7032. 10.1038/s41598-018-25226-w

Bochereau, S., Dzidek, B., Adams, M., Hayward, V., 2017. Characterizing and Imaging Gross and Real Finger Contacts under Dynamic Loading. IEEE Trans. Haptics 10, 456–465. 10.1109/TOH.2017.2686849

Brinkerhoff, S.A., Mathew, G.M., Murrah, W.M., Chang, A.-M., Roper, J.A., Neely, K.A., 2022. Sleep restriction impairs visually and memory-guided force control. PLOS ONE 17, e0274121. 10.1371/journal.pone.0274121

Coombes, S.A., Corcos, D.M., Vaillancourt, D.E., 2011. Spatiotemporal tuning of brain activity and force performance. NeuroImage 54, 2226–2236. 10.1016/j.neuroimage.2010.10.003

Cuadra, C., Latash, M.L., 2019. Exploring the Concept of Iso-perceptual Manifold (IPM): A Study of Finger Force-Matching Tasks. Neuroscience 401, 130–141. 10.1016/j.neuroscience.2019.01.016

Delhaye, B.P., Jarocka, E., Barrea, A., Thonnard, J.-L., Edin, B., Lefèvre, P., 2021a. High-resolution imaging of skin deformation shows that afferents from human fingertips signal slip onset. eLife 10, e64679. 10.7554/eLife.64679

Delhaye, B.P., Schiltz, F., Barrea, A., Thonnard, J.-L., Lefèvre, P., 2021b. Measuring fingerpad deformation during active object manipulation. J. Neurophysiol. 126, 1455–1464. 10.1152/jn.00358.2021

Diedrichsen, J., Shadmehr, R., Ivry, R.B., 2010. The coordination of movement: optimal feedback control and beyond. Trends Cogn. Sci. 14, 31–9. 10.1016/j.tics.2009.11.004

Dzidek, B., Bochereau, S., Johnson, S., Hayward, V., Adams, M., 2016. Frictional dynamics of finger pads are governed by four length-scales and two time-scales, in: 2016 IEEE Haptics Symposium (HAPTICS). Presented at the 2016 IEEE Haptics Symposium (HAPTICS), IEEE, Philadelphia, PA, pp. 161–166. 10.1109/HAPTICS.2016.7463171

Dzidek, B., Bochereau, S., Johnson, S.A., Hayward, V., Adams, M.J., 2017. Why pens have rubbery grips. Proc. Natl. Acad. Sci. 114, 10864–10869. 10.1073/pnas.1706233114

Dzidek, B.M., Adams, M., Zhang, Z., Johnson, S., Bochereau, S., Hayward, V., 2014. Role of Occlusion in Non-Coulombic Slip of the Finger Pad, in: Auvray, M., Duriez, C. (Eds.), Haptics: Neuroscience, Devices, Modeling, and Applications,Lecture Notes in Computer Science. Springer Berlin Heidelberg, Berlin, Heidelberg, pp. 109–116. 10.1007/978-3-662-44193-0_15

JASP Team, 2024. JASP (Version 0.18.3)[Computer software].

Jo, H.J., Ambike, S., Lewis, M.M., Huang, X., Latash, M.L., 2016. Finger force changes in the absence of visual feedback in patients with Parkinson’s disease. Clin. Neurophysiol. 127, 684–692. 10.1016/j.clinph.2015.05.023

Johansson, R.S., Westling, G., 1988. Programmed and triggered actions to rapid load changes during precision grip. Exp. Brain Res. 71, 72–86. 10.1007/BF00247523

Johansson, R.S., Westling, G., 1987. Signals in tactile afferents from the fingers eliciting adaptive motor responses during precision grip. Exp. Brain Res. 66. 10.1007/BF00236210

Johansson, R.S., Westling, G., 1984. Roles of glabrous skin receptors and sensorimotor memory in automatic control of precision grip when lifting rougher or more slippery objects. Exp. Brain Res. 56. 10.1007/BF00237997

Naik, A., Ambike, S., 2022. Expectation of volitional arm movement has prolonged effects on the grip force exerted on a pinched object. Exp. Brain Res. 240, 2607–2621. 10.1007/s00221-022-06438-z

Neely, K.A., Mohanty, S., Schmitt, L.M., Wang, Z., Sweeney, J.A., Mosconi, M.W., 2019. Motor Memory Deficits Contribute to Motor Impairments in Autism Spectrum Disorder. J. Autism Dev. Disord. 49, 2675–2684. 10.1007/s10803-016-2806-5

Olczak, D., Sukumar, V., Pruszynski, J.A., 2018. Edge orientation perception during active touch. J. Neurophysiol. 120, 2423–2429. 10.1152/jn.00280.2018

Pasumarty, S.M., Johnson, S.A., Watson, S.A., Adams, M.J., 2011. Friction of the Human Finger Pad: Influence of Moisture, Occlusion and Velocity. Tribol. Lett. 44, 117–137. 10.1007/s11249-011-9828-0

Peters, R.M., Hackeman, E., Goldreich, D., 2009. Diminutive Digits Discern Delicate Details: Fingertip Size and the Sex Difference in Tactile Spatial Acuity. J. Neurosci. 29, 15756–15761. 10.1523/JNEUROSCI.3684-09.2009

Poon, C., Chin-Cottongim, L.G., Coombes, S.A., Corcos, D.M., Vaillancourt, D.E., 2012. Spatiotemporal dynamics of brain activity during the transition from visually guided to memory-guided force control. J. Neurophysiol. 108, 1335–1348. 10.1152/jn.00972.2011

Reinkensmeyer, D.J., Akoner, O.M., Ferris, D.P., Gordon, K.E., 2009. Slacking by the human motor system: Computational models and implications for robotic orthoses, in: 2009 Annual International Conference of the IEEE Engineering in Medicine and Biology Society. Presented at the 2009 Annual International Conference of the IEEE Engineering in Medicine and Biology Society, IEEE, Minneapolis, MN, pp. 2129–2132. 10.1109/IEMBS.2009.5333978

Reschechtko, S., Cuadra, C., Latash, M.L., 2017a. Force illusions and drifts observed during muscle vibration. J. Neurophysiol. jn.00563.2017. 10.1152/jn.00563.2017

Reschechtko, S., Pangan, W.R., Zadeh, R.Z., Pruszynski, J.A., 2024. Single-shot detection of microscale tactile features. 10.1101/2024.09.03.610877

Reschechtko, S., Zatsiorsky, V.M., Latash, M.L., 2017b. The synergic control of multi-finger force production: stability of explicit and implicit task components. Exp. Brain Res. 235, 1–14. 10.1007/s00221-016-4768-4

Scholz, J.P., Schöner, G., Latash, M.L., 2000. Identifying the control structure of multijoint coordination during pistol shooting. Exp. Brain Res. 135, 382–404. 10.1007/s002210000540

Smith, B.W., Rowe, J.B., Reinkensmeyer, D.J., 2018. Real-time slacking as a default mode of grip force control: implications for force minimization and personal grip force variation. J. Neurophysiol. 120, 2107–2120. 10.1152/jn.00700.2017

Todorov, E., Jordan, M.I., 2002. Optimal feedback control as a theory of motor coordination. Nat. Neurosci. 5, 1226–1235. 10.1038/nn963

Vaillancourt, D.E., Russell, D.M., 2002. Temporal capacity of short-term visuomotor memory in continuous force production. Exp. Brain Res. 145, 275–285. 10.1007/s00221-002-1081-1

Vaillancourt, D.E., Slifkin, A.B., Newell, K.M., 2001. Visual control of isometric force in Parkinson’s disease. Neuropsychologia 39, 1410–1418. 10.1016/S0028-3932(01)00061-6

Vaillancourt, D.E., Thulborn, K.R., Corcos, D.M., 2003. Neural Basis for the Processes That Underlie Visually Guided and Internally Guided Force Control in Humans. J. Neurophysiol. 90, 3330–3340. 10.1152/jn.00394.2003

Vallat, R., 2018. Pingouin: statistics in Python. J. Open Source Softw. 3, 1026. 10.21105/joss.01026

